# *In situ* structure and rotary states of mitochondrial ATP synthase in whole cells

**DOI:** 10.1101/2024.03.27.586927

**Authors:** Lea Dietrich, Ahmed-Noor Adam Agip, Andre Schwarz, Christina Kunz, Werner Kühlbrandt

## Abstract

Cells depend on a continuous supply of ATP, the universal energy currency. In mitochondria, ATP is produced by a series of redox reactions, whereby an electrochemical gradient is established across the inner mitochondrial membrane. The ATP synthase harnesses the energy of the gradient to generate ATP from ADP and inorganic phosphate. We determined the structure of ATP synthase within mitochondria of the unicellular alga *Polytomella* by electron cryo-tomography. Sub-tomogram averaging revealed six rotary positions of the central stalk, subclassified into 21 substates of the F_1_ head. The *Polytomella* ATP synthase forms helical arrays with multiple adjacent rows defining the cristae ridges. The structure of ATP synthase under native operating conditions in the presence of a membrane potential represents a pivotal step toward the analysis of membrane protein complexes *in situ*.

## Introduction

The F_1_F_o_ ATP synthase plays a crucial role in biological energy conversion in mitochondria, bacteria and chloroplasts. In mitochondrial oxidative phosphorylation, exergonic electron transfer through respiratory chain enzymes and mobile electron carriers in the inner mitochondrial membrane (IMM) and the intermembrane space is coupled to endergonic proton translocation across the IMM, establishing an electrochemical proton gradient. The energy of the proton gradient is harnessed by the ATP synthase to generate ATP. Working like a molecular turbine, the ATP synthase produces adenosine triphosphate (ATP) from ADP and inorganic phosphate by rotary catalysis. The ATP thus generated is the primary energy source for virtually all cellular processes.

The structure of mitochondrial ATP synthase has been investigated by electron microscopy for more than six decades (*1, 2*). The complex consists of the water-soluble spherical F_1_ head (*3*) and the membrane-spanning F_o_ subcomplex (*4*). Under physiological conditions, the F_o_ subcomplex generates the torque that powers ATP production in the F_1_ head (*5*). A ring of 8 to 17 hydrophobic c-subunits (*6–10*) undergoes a complete rotation, transferring protons from the intermembrane space (pH 6.8-7.4) (*11, 12*) to the matrix (∼pH 8) (*13, 14*). The energy derived from downhill proton translocation propels the rotation of the *c*-ring and the firmly attached central stalk of subunits γ, δ, ε, transmitting the torque generated by F_o_ to drive ATP synthesis in the F_1_ head. The rotation of the central stalk induces conformational changes via the two-helix bundle of the γ subunit. The peripheral stalk establishes a second connection between the F_1_ and F_o_ subcomplexes, acting as a stator and preventing idle rotation of the F_1_ head during catalysis.

Detailed structures of the soluble F_1_ part were obtained by X-ray crystallography, but structures of the whole, highly dynamic F_1_F_o_ ATP synthase had to await high-resolution electron cryo-microscopy (cryoEM), which has delivered structures of the intact mitochondrial complex at resolutions ranging from 2.8 Å (*15*) to 3.8 Å (*9, 16, 17*). Unlike bacterial and chloroplast ATP synthases, the mitochondrial ATP synthase of all species investigated so far forms dimers in the IMM (*18*), creating the mitochondrial cristae which are thought to work as local proton reservoirs (*19*).

F-type ATP synthases are subdivided into four types (*19*). In type I, found in mammals and fungi, the dimer angle between individual monomers is 86° ((EMD-7067) (*20*), (EMD-11436) (*21*)), Type-II ATP synthase dimers found in the unicellular green algae *Polytomella sp*. and *Chlamydomonas reinhardtii* are characterized by a bulky, rigid peripheral stalk composed of 10 ATP synthase-associated (ASA) subunits (*22, 23*). The reported dimer angle was 56°. In the IMM, the dimers assemble into long rows that determine the cristae architecture (*24*). Type IV, like I & II is V-shaped with a dimer angle of 56° (*25*). In type III, the dimer angle is 0° and tubular cristae result from dimers assembling into long helical ribbons (*26*)

Electron cryo-tomography (cryo-ET) of mitochondrial membranes revealed that the rows of type-I dimers extend to more than 1 μm in length in bovine, yeast and green plant mitochondria (*18, 27*). Loss of the interface subunits *g* and *e* in *S. cerevisiae* rendered the protein monomeric, resulting in balloon-shaped cristae with a random distribution of monomers in the membrane (*28*). Conversely, isolated, detergent-solubilized ATP synthase dimers reconstituted into proteoliposomes demonstrated spontaneous self-assembly into rows that induce local membrane curvature, with geometries closely resembling those observed in the IMM (*24*).

For a comprehensive understanding of how ATP synthase works, it is essential to investigate its structure and mechanism *in situ*, ensuring that its native environment, in particular the proton motive force (pmf) that drives ATP synthesis, is preserved. We applied cryo-ET and subtomogram averaging (StA) of focussed ion beam-milled (FIB-milled) *Polytomella* cells to image the mitochondrial ATP synthase *in situ*. The *Polytomella* ATP synthase is well-suited to this approach because of its stability and distinctive shape, and because mitochondria account for a large fraction of the cell volume.

## Results and Discussion

### *In situ* structure of the mitochondrial ATP synthase

To determine the *in situ* structure and spatial arrangement of the F-type ATP synthase within mitochondria, *Polytomella* cells in the growth phase were flash-frozen on EM grids, followed by FIB-milling of cryo-lamellae and data acquisition for cryo-ET. The thickness of the cryo-lamellae ranged from to ∼60 - 220 nm, as required for high resolution cryo-ET (Fig.S1). A total of 552 tomograms were acquired, of which 255 were used for structure determination. Tomograms were selected on the basis of lamella thickness, tilt series alignment and a metric score from cryo-ET software data processing (see Materials and Methods).

We manually selected 1,399 coordinates of ATP synthase particles to provide a reference for template matching (*29*). Briefly, ATP synthase monomers were identified by their characteristic mushroom shape, protruding ∼15 nm from the membrane. The bulky peripheral stalk is clearly visible in top, side and front views of individual dimers (Fig.S2). 3D reconstruction resulted in an initial low-resolution volume of ATP synthase, which was used as input for template matching (*29*). Initial particle extraction resulted in 363,061 initial particles. False positives were removed through iterative rounds of classification with alignment in Relion (*30*). 3D refinement of the initial dimer data set with C2 symmetry resulted in a consensus map at ∼17Å resolution (Fig.S2). The initial subtomogram average was subjected again to template matching (*29*) to enhance the particle number and accuracy of picks. The final round of classification yielded 131,411 particles contributing to the consensus map. Refinement using M (*31*) with C2 symmetry imposed resulted in the final consensus map at a local resolution up to 4.2 Å (Fig.1).

**Fig. 1.**
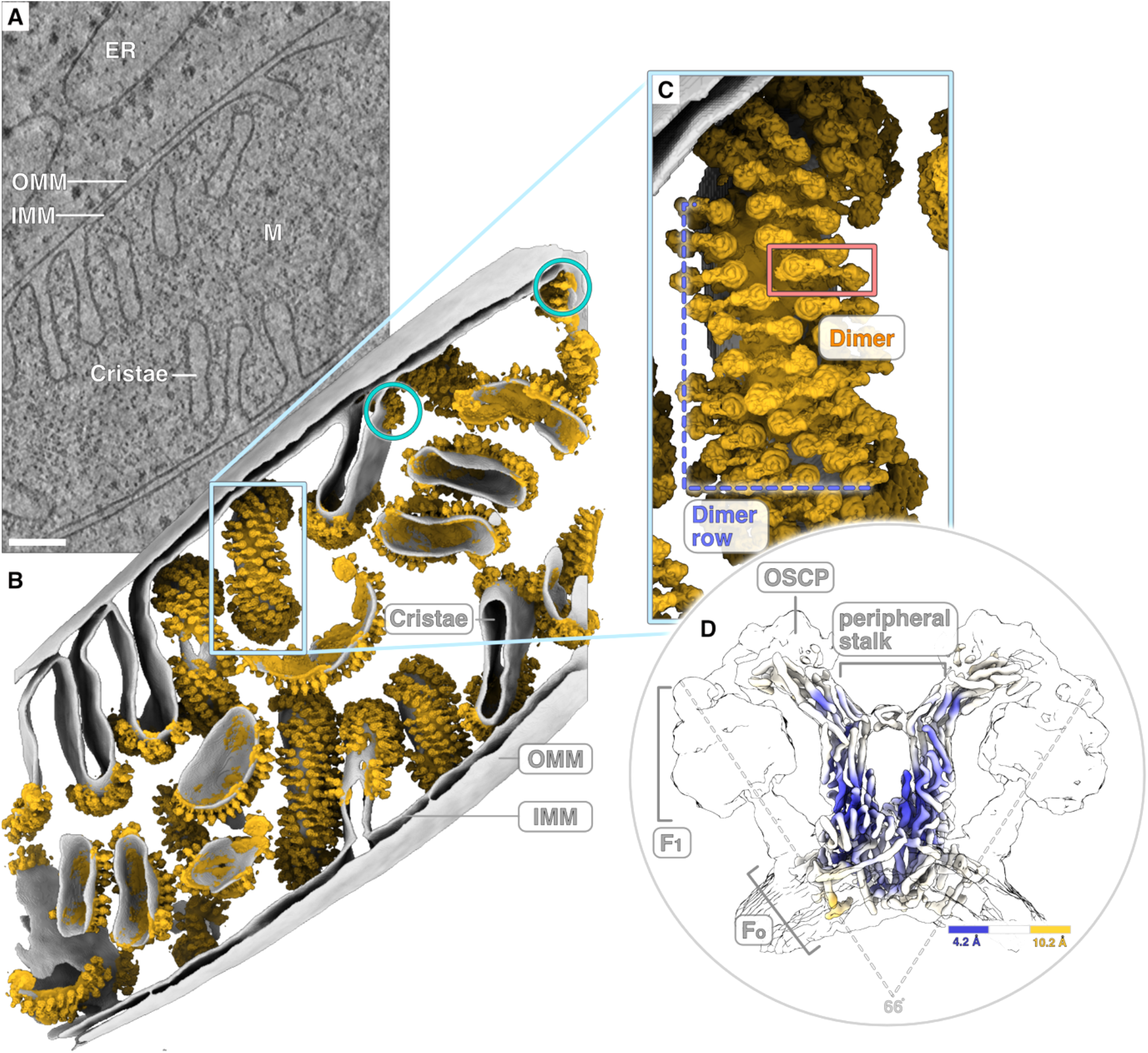
*In situ* structure of the mitochondrial ATP synthase. **(A**) Tomographic slice showing a mitochondrion of the green algae *Polytomella sp*. next to the endoplasmatic reticulum (ER). The inner and outer mitochondrial membranes (IMM and OMM) are clearly visible, as are the mitochondrial cristae protruding into the matrix (M). Scale bar: 100 nm. (**B/C)** The 3D volume of a mitochondrion reveals densely packed ATP synthase dimers covering the cristae ridges. ATP synthase dimers are arranged in a near-helical pattern with parallel multiple rows. While predominantly located at the cristae ridges, ATP synthases are also found close to the cristae junctions (cyan circles). (**D)** Local resolution of the ATP synthase consensus map indicates 4.2 Å resolution at the core of the bulky peripheral stalk. The F_1_ and F_o_ subcomplexes are not as well resolved because they are cylindrically averaged due to rotation around their axes. ATP synthase monomers include an angle of 66° which induces membrane bending.

### ATP synthase dimer rows form higher-order assemblies creating the cristae ridges

To elucidate the arrangement of ATP synthase dimers in the inner mitochondrial membrane we combined the poses of the consensus refinement with membrane segmentation. Mapping back the coordinates of the consensus map confirmed previous reports of dimer rows along the cristae ridges (*28, 32*). In addition, we see that multiple dimer rows run next to each other in a regular, helical pattern, twisting around the cristae ridges and stabilizing the disk-like cristae (Fig.1). The dimer rows follow a left-handed helical geometry, deviating from a perfect helix due to flexible inter-dimer packing (see Movie S1). No additional protein density was detected at the interface that might define the spacing (Fig.2B).

**Fig. 2.**
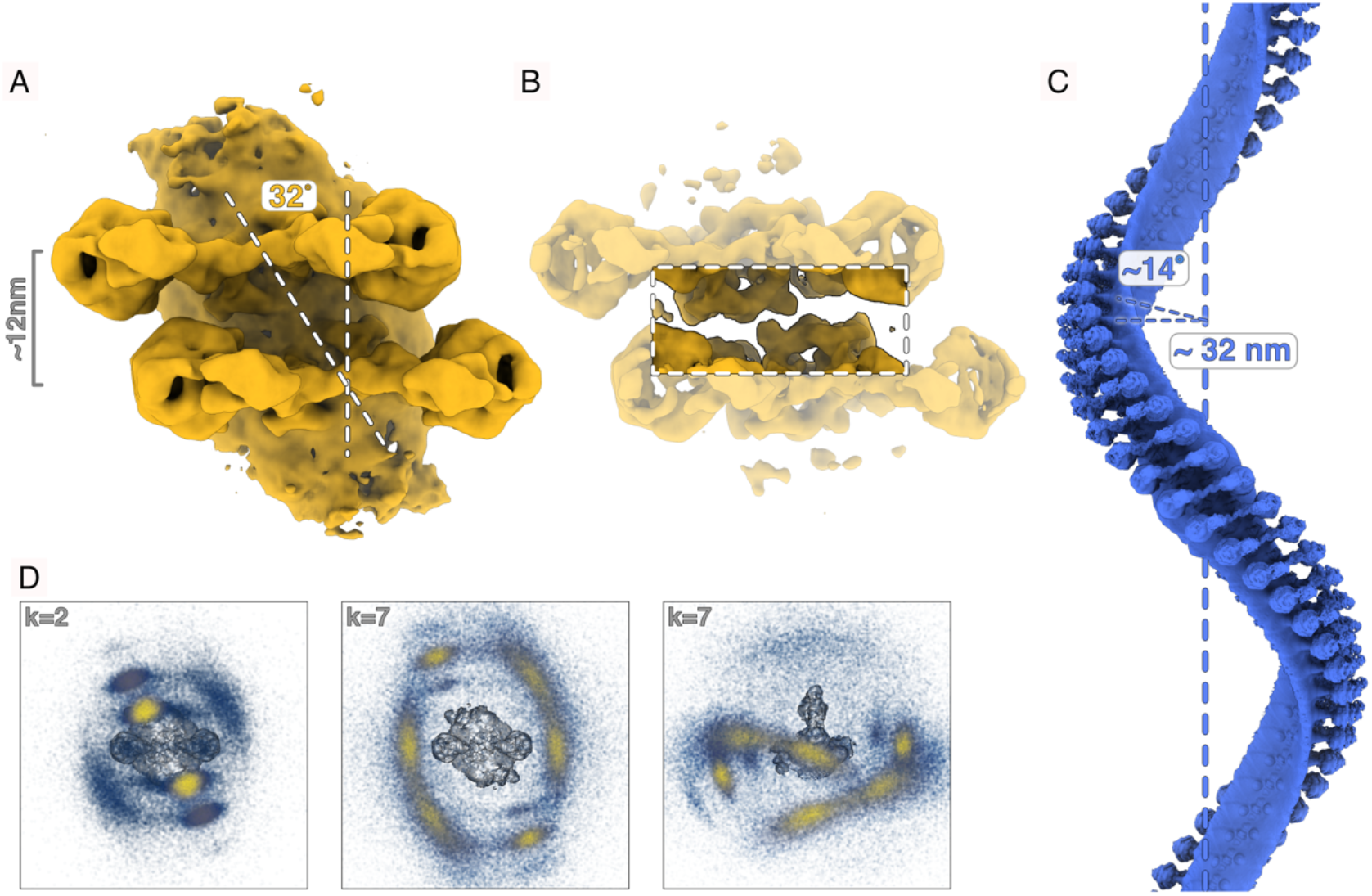
ATP synthase tetramers are defined as the repeating units of dimer rows. **(A)** Within each ATP synthase tetramer, dimers are separated by about 12 nm with a 32° offset. **(B)** At a lower isosurface threshold a hook-like structure becomes apparent which may play a role in the spacing of ATP synthase dimers along the rows. (**C)** The model of a perfect helix of *Polytomella* ATP synthase dimers indicates a helix radius of 32 nm and a helical twist of approx.14°. (**D)** Nearest-neighbour analysis indicates the spatial distribution of the ATP synthase *in situ*. k is the number of adjacent particles in the search, including itself. k=2 gives the average distance between each particle position and its first nearest neighbour, confirming an approximately uniform spacing within the dimer rows. k=7 (central and right panels showing the top and side view of a dimer in a row) represents the average distance between each particle position and its six nearest neighbours and indicates the regular spacing between adjacent rows. The consensus map of ATP synthase is shown as a reference point in the centre of each panel. Particle positions are coloured by local density (yellow = high local density).

ATP synthase dimer ribbons are mostly confined to the cristae ridges (*24, 28*), but can extend to the cristae junctions (Figure 1: cyan circles). The cristae ridges are stabilized by at least two dimer rows defining the cristae shape. Nearest-neighbour coordinates for neighbouring rows indicated a consistent spacing of adjacent rows with an average distance of 29 nm. The mean distance between the centres of adjacent dimers in a row is ∼12 nm ± 2 nm (Fig.2).

Our previous work on purified mitochondria and isolated crista vesicles demonstrated the ability of ATP synthase dimers to self-assemble into rows (*24*). Cryo-ET of mitochondria isolated from *Polytomella* indicated a similar stacking geometry. We now extend this knowledge by showing that the type-II ATP synthase of green algae forms left-handed helical arrays deviating from right-handed helices formed by type-III ATP synthases found in ciliates. The rows run parallel to each other, maximizing coverage of the surface area along the cristae ridges. The respiratory chain complexes are most likely confined to the flat membrane regions on either side of the dimer rows.

### Atomic model of the ATP synthase dimer *in situ*

The local resolution of the subtomogram average map varies, with the highest resolution (4.2 Å) found in the peripheral stalk at the dimer centre. The peripheral stalk subunits ASA1-10, mostly consisting of tightly packed alpha-helices, contribute significantly to the stability of the *Polytomella* ATP synthase dimer (*22, 33*).

Starting from the 2.8 Å single-particle model (PDB: 6rd4), all ASA subunits were docked into the StA density. The high local resolution of the peripheral stalk enabled us to trace with confidence the alpha carbon chain of subunits ASA1, ASA3, ASA5, ASA6, ASA7, ASA8, ASA9, ASA10, and the proton-translocating subunit *a*. Densities of bulky aromatic sidechains served as guide points to ensure the correct sequence register (Fig.S6). We were able to fit 89.4% of the peripheral stalk subunits, including subunit *a*, which compares well to the previous 90.8% fit to the single-particle 2.8 Å map (Table S1).

Our fit follows the amino acid sequence positions along the polypeptide chain of the membrane-embedded stator, subunit *a*, which forms a conserved near-horizontal helix hairpin in the middle of the membrane (*34*). The density of subunit *a* allowed us to model the loop of 13 amino acids between H5 and H6 (Fig.3C) that was missing in the single-particle map. In this map (*22, 23, 34*), ASA3 forms a characteristic Armadillo repeat on the matrix side of the membrane. Surprisingly, we found a major new density continuous with ASA3 that forms a loop and extends horizontally along the membrane surface. As the AlphaFold (*35*) prediction for the *Polytomella* ATP synthase was inaccurate in this region, we first built the ASA3 extension based on a homology model from the related *Chlamydomonas reinhardtii*. We then applied flexible fitting (*36*), and mutated the sequence to that of *Polytomella*. The ASA3 extension is modelled mostly as α-helical, followed by a transmembrane α-helix at the N-terminus (Fig.3).

**Fig. 3.**
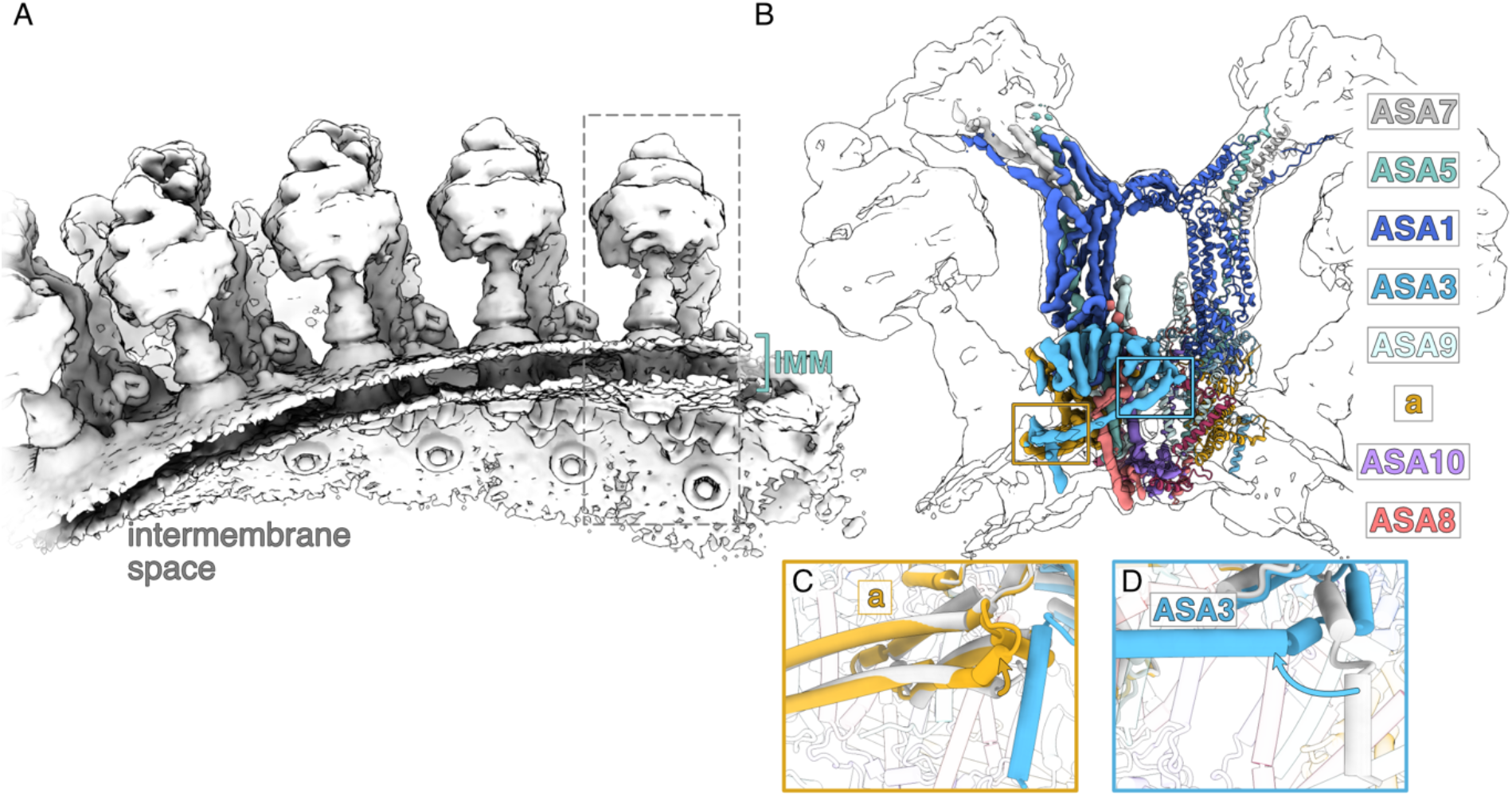
Model of ATP synthase dimers *in situ*. **(A)** Closeup of an ATP synthase dimer row, seen from the side. **(B)** Model of the peripheral stalk of ATP synthase *in situ*, based on the model of our 2.8 Å single particle structure (PDB: 6rd4) as a starting point and refined against the consensus map. The resolution of most peripheral stalk subunits was sufficient to dock the model into our map with confidence. Eight out of the 10 type-II-specific ASA subunits and the highly conserved subunit *a* were assigned. **(C/D)** Major changes between the published (white) and *in situ* model (coloured) were found in the subunits *a* and ASA3.

On the basis of this model, we propose that the ASA3 N-terminus starts as a transmembrane helix adjacent to the quasi-horizontal membrane-intrinsic helix hairpin of subunit *a*. ASA3 runs along the membrane surface for a stretch of 50 residues. Together with ASA9, the loop connecting the new density to the Armadillo turn of ASA3 forms a cleft in close proximity to the adjacent dimer. While we do not observe a clear inter-dimer contact in our tetramer map (Fig.2), the hook-shaped cleft formed by ASA3 and ASA9 may play a role in row formation by helping to interlock the dimers and thus maintain a quasi-regular spacing. Previous single particle analysis of the porcine ATP synthase tetramer described four sites participating in tetramer formation (*37*). The structure was solved in presence of the inhibitory protein 1 (IF1) bridging the F_1_ heads of neighbouring dimers. There is no known homologue of IF1 in *Polytomella*, and we do not observe a density reminiscent of such a protein.

The extra elongated density of subunit ASA3, which is not present in the single-particle cryoEM structure, is most likely due to the enhanced stability of the complex in the membrane lipid environment. The long N-terminal helix of ASA3 may be important for stabilizing the loop between the intramembrane horizontal helix pair (consisting of helices 5 and 6) of the stator subunit *a* which is likewise unstructured in the detergent-solubilized dimer (*22, 33*). The functional implications, in particular how the presence or absence of this loop in detergent or lipid nanodiscs affects ATP synthase activity or regulation, are currently unknown.

Comparison with mitochondrial ATP synthases from other species reveals that, although ASA3 is alga-specific, its absence in mammals and fungi is compensated by structurally homologous proteins, which play the same role in stabilizing the subunit *a* loop (Fig.S4). In bovine ATP synthase, the loop between H5 and H6 of subunit *a* interacts with the diabetes-associated protein in insulin-sensitive tissue (DAPIT) (Fig. S4B) (*17*), indicating a likely medical significance.

### Classification of rotary states in ATP synthesis mode

The mitochondrial ATP synthase operates by rotary catalysis, powered by the electrochemical proton gradient across the IMM, which induces multiple conformational changes. To identify distinct catalytically-relevant states, we applied symmetry expansion to the subtomogram average of the dimer, followed by focused classification, masking the central stalk of the monomer without imposing C2 symmetry (see materials and methods and Fig.S3). In this way we were able to distinguish six principal rotary states, which were further refined to resolve distinct features. The resolution of the resulting maps ranges from ∼10-13 Å, reflecting the limited number of particles in each class (Fig.4A). Visual inspection and superposition of classes confirmed distinct relative positions of the central and peripheral stalks. The principal state that most closely resembles the primary rotary state 1 identified in the single-particle map (PDB:6rd9, (*22*)), we refer to as principal state I. The other, less populated rotary states were put into a sequence that reflects the counter-clockwise rotation of the central stalk as seen from the matrix (Fig.S5). To separate rotary substates capturing the movement in the F_1_ head, we applied a second focused classification to the F_1_-F_o_ region of the monomer. The classification revealed two to four rotary substates within each principal state that were identified with confidence. Their superposition (Fig.4C) shows a rotation of the F_1_ head around the axis of the central stalk between the first and later sub-states. In addition, slight differences in the position of the flexible OSCP joint were resolved between some of the classes. These differences most likely result from the hinge motion of the OSCP which ensures flexible coupling of the F_1_ and the F_o_ motors (*22*).

**Fig. 4.**
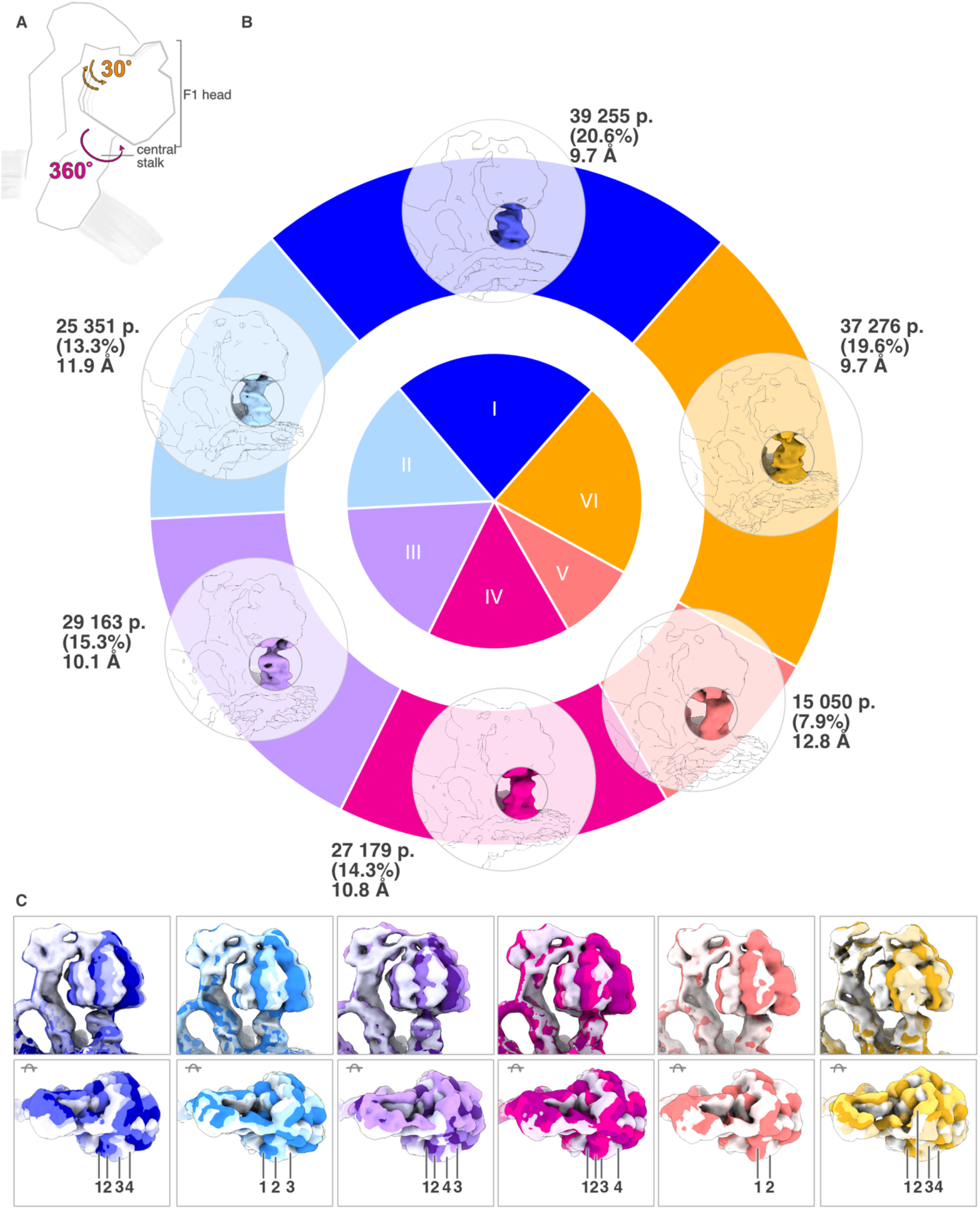
Rotational states of ATP synthase resolved *in situ*. **(A)** Schematic representation of the two rotary modes that we were able to separate. **(B)** The dominant motion defining the principal rotary states is the 360° rotation of the central stalk. This rotation was resolved with a focused mask around the central stalk and yielded the six principal states. **(C)** The second, more subtle movement is that of the F_1_ head, which moves by about 30° together with the central stalk and then snaps back (*15*). We resolved this movement with a mask around the F_1_F_o_ part of the ATP synthase, resulting in 2-4 rotational substates per principal state, whereby the F_1_ head rotates around the axis of the central stalk.

In total, 91% of the initial particles used for classification were assigned to a principal rotary state and 75% of the subtomograms were assigned to a rotary substate.

As with our previous single-particle 2.8 Å map (*22*), we performed focused classification and refinement to dissect the different rotary states of the ATP synthase, making use of symmetry expansion to increase the number of particles and improve the map resolution. The presence of a membrane potential in our system, leading to continuous rotation of the ATP synthases, enabled us to determine several additional substates that are energetically favourable and have not been observed before. The prevalence of six specific conformations indicates that energy minima exist during turnover. However, the moderate resolution of the rotary states also indicates a considerable degree of heterogeneity within these classes.

*In situ* structural biology is proving to be a powerful tool for unravelling both major and minor movements during the catalytic cycle of ATP synthase. Furthermore, these catalytically relevant states are observed in the cell, in the presence of a membrane potential and in the absence of detergents, inhibitors or any pharmacological agents (*38*). We present the first *in situ* structures of the complex under native conditions during ATP synthesis, which have been difficult to access in the field of bioenergetics.

Together with the central stalk, the *c*-ring of the *Polytomella* ATP synthase undergoes a 360° rotation, during which 10 protons are translocated across the inner mitochondrial membrane. The outer transmembrane helices of the 10 *c*-subunits in the ring are resolved in the consensus map and are separated by ∼36°, as expected (Fig.5A-C). The clear density in the centre of the *c*-ring is most likely due to two or more lipids of the two membrane leaflets arranged end-to-end. In single-particle maps of bovine and ovine ATP synthase, similar densities have been found in the same position (*9, 21*). The highly hydrophobic internal space of the *c*-ring is most likely occupied by lipids at an early stage of complex assembly.

**Fig. 5.**
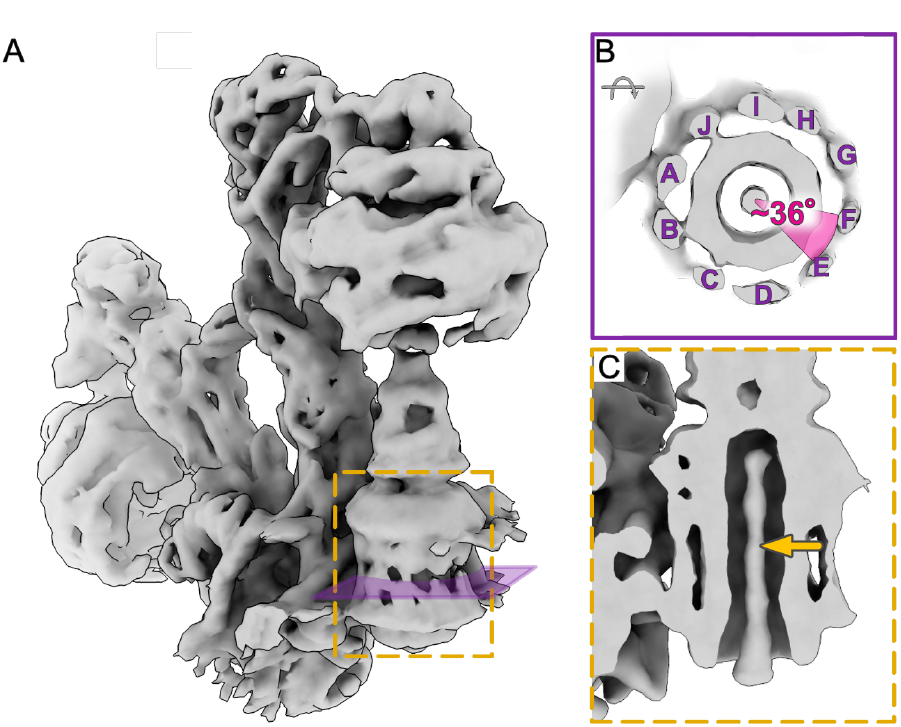
Lipid density and c-ring of ATP synthase. **(A)** Consensus map of the *Polytomella* ATP synthase showing the *c*-ring (dashed yellow rectangle). (**B)** Cross section through the c-ring density revealing the outer α-helices of the helix hairpins of the 10 *c*-subunits in the ring, separated by ∼36°. (**C)** Density in the central space of the *c*-ring is most probably due to two or more membrane lipids arranged end-to-end (arrow).

## Summary

We present our structure of ATP synthase dimers within cells of the model organism *Polytomella sp*. determined by *in situ* electron cryo-tomography and subtomogram averaging to a resolution of up to 4.2 Å. We find that ATP synthases of this type form higher-order assemblies, describing a left-handed helix wrapping around the ridges of the inner membrane cristae. Our map reveals an additional, previously unknown helix density of subunit ASA3. This subunit plays an important role in stabilizing the conserved loop of the proton-translocating stator subunit *a* that is essential for ATP synthesis. Our study demonstrates the power of *in situ* structural biology that preserves interactions which may otherwise be lost by protein isolation and purification. We show that is possible to dissect the various rotary states of the ATP synthase within mitochondria of whole, flash-frozen cells. A complete picture of oxidative phosphorylation requires not only precise knowledge of the distribution and arrangement of ATP synthases in the cell, in relation to other complexes in molecular bioenergetics. Our work provides an essential foundation on the analysis of *in situ* structures not only for bioenergetic complexes, but of protein assemblies in the context of their native membrane, and, more generally, in their cellular environment.

## Acknowledgments

We thank the Central Electron Microscopy Facility of the Max Planck Institute of Biophysics lead by Sonja Welsch for cryo-EM infrastructure and technical support as well as help with the manuscript. Juan Castillo-Hernández, Özkan Yildiz, the Central IT team, and the Max Planck Computing and Data Facility for support in cryo-EM data processing. Many thanks to Beata Turoňová for data processing support and Bonnie Murphy and Niklas Klusch for their expertise and discussion. We thank Janet Vonck for tremendous help with model building and Pamela Ornelas for discussion. We thank Tristan Coll for advice on model building.

## Funding

Max Planck Society

Deutsche Forschungsgemeinschaft (FOR2848)

## Author contributions

Project conceptualization and initiation: L.D., WK

Sample preparation and lamella milling: L.D., C.K.

Data collection and processing: L.D.

Data analysis: L.D., A.S.

Model building: L.D., A.-N.A.A.

Writing: L.D., A.-N.A.A., W.K.

## Competing interests

Authors declare that they have no competing interests.

## Data and materials availability

Full dimer map: EMD-XXXX

Local refinement map of the peripheral stalk: EMD-XXXX

Density-modified map of peripheral stalk with model: EMD-XXXX

## Supplementary Materials

Materials and Methods

Figs. S1 to S6

Table S1

References

Movie S1

## Notes

### Competing Interest Statement

The authors have declared no competing interest.

