## Supplementary material for "*In situ* structure and rotary states of mitochondrial ATP synthase in whole cells": Supplemetary

### Materials and Methods

#### Sample Preparation

*Polytomella sp.* was cultivated at room temperature without agitation, following previously described methods (Allegretti et al., 2015). A 50 ml culture was inoculated with 50 µl of a stock culture and allowed to grow overnight until reaching an optical density ($OD_{600}$) of 0.8-1. Subsequently, 3 µl of the culture were applied to an EM grid (Quantifoil #200 R2/2) and blotted from the back side using an EM GP2 plunger (Leica) before freezing in liquid ethane. Cell activity was verified under a light microscope prior to freezing.

#### Focused Ion Beam-Scanning Electron Microscopy

Grids were clipped into modified Autogrids for low-angle FIB milling. The clipped grids were loaded into an Aquilos cryo-FIB (Thermo Scientific) scanning electron microscope for lamella preparation, following the procedure described by Wagner *et al.* (*1*). In brief, loaded grids were coated with organometallic platinum using the gas injection system (GIS), followed by a layer of inorganic platinum applied via sputter coating. Single cells were identified as suitable positions for lamella preparation. Eucentricity was adjusted manually for each position and the milling angle was set to 8°. Lamella preparation was performed in a step-wise process, starting from 5 µm, then to 3 µm, and finally to 1 µm lamella thickness, using gallium beam milling currents of 1 nA, 0.5 nA, and 0.3 nA, respectively. For the final polishing, 30-50 pA was used to achieve a lamella thickness of <250 nm (see Fig.S1). After polishing, the lamellae were coated with an additional thin layer of inorganic platinum.

#### Electron Cryo-Tomography Data collection

Cryo-electron tomography (Cryo-ET) data were acquired using a Titan Krios G2 transmission electron microscope (Thermo Scientific), operated at 300keV in EFTEM mode and equipped with a BioQuantum-K3 imaging filter (Gatan). To localize mitochondria and define areas suitable for tilt series collection, overview montages of individual lamellae were acquired at a nominal magnification of 6,500x. Tilt series were recorded with SerialEM (version 4.0.1) (*2*) at a nominal magnification of 53,000x, corresponding to a calibrated pixel size of 1.63 Å/pixel. We applied a dose-symmetric tilt scheme (*3*) , starting with an 8° pretilt and proceeding with 2° increments. The dose per tilt angle was set at 1.9-2 e/Å$^2$, adding up to a total dose of 115-120 e/Å$^2$ per tilt series. The energy filter slit width was set to 20 eV, and the set defocus per tilt series cycled between -2.5 and -4.5 µm. Dose-fractionated frames were aligned using on-the-fly motion correction in SerialEM.

#### Data Processing:

Visual inspection was used to clean the tilt series by eliminating dark, blurred, or off-target images before proceeding with tilt series alignment and tomogram reconstruction using AreTomo (version 1.3.0) (*4*). Tomograms binned to a pixel size of 13.5 Å/pixel were used for initial quality assessment, focusing on tilt series alignment and tomogram thickness. For subsequent data processing with Relion-3.1 (*5*) and M (*6*), tomograms were reconstructed in Warp (*7*) using the alignment parameters obtained from AreTomo. At this stage, the data set was cleaned further, removing tilt images with low resolution and/or astigmatism.

Template matching was conducted using the STOPGAP toolbox (*8*). An initial reference was generated from 1,399 manually picked particles, employing STOPGAP subtomogram averaging at a pixel size of 9.78 Å/pixel. The reference was sampled across each tomogram at an angular sampling rate of 30°. Peak extraction was performed by opening each score volume in Chimera (*9*) and inspecting it in solid mode. The threshold for peak extraction for each score volume was determined individually, guided by prior knowledge of localization and row formation to discern true positive peaks. The resulting STOPGAP star file was converted into a Warp-compatible star file using the dynamo2m toolbox (https://github.com/alisterburt/dynamo2m). Subtomograms were extracted in Warp (*7*) and subjected to two rounds of 3D classification in Relion-3.1 resulting in an initial consensus map at 17 Å resolution from 38,970 subtomograms. A second iteration of template matching was performed using the generated map as new reference, resulting in 238,622 identified positions.

Through iterative rounds of 3D classification in Relion-3.1 a consensus map at a resolution of ~15 Å at a pixel size of 6.8 Å/pixel was obtained. Particle positions were refined in M and particles extracted at a pixel size of 3.4 Å/pixel. The initial consensus map was refined in Relion- 3.1 without symmetry and with a mask around the whole dimer and subsequently centred along the symmetry axis and refinement in M with C2 symmetry imposed, resulting in a map of the whole dimer with 8.4 Å global resolution. Focusing on the rigid peripheral stalk, the global resolution was further improved to 6.8 Å and local resolution to 4.2 Å.

**Classification of the rotary states**
In order to dissect the different rotary states of the $F_1F_o$ subcomplex, symmetry expansion was applied after refinement of the whole dimer in C2 at a pixel size of 3.4 Å/pixel. Subsequently the particles were aligned on the peripheral stalk of one monomer in C1. Differentiation of principal states was achieved by 3D classification without alignment focusing on the central stalk while excluding the $F_1$-head and the membrane. We asked for seven classes of which six resulted in a refined map. To separate further substates of the $F_1$ head, which is known to rotate by about 30° along the central stalk (*10*), a mask comprising the $F_1F_o$ part was applied. Again, we asked for 7 classes from which only 2-4 were subjected to further refinement based on particle number and map quality. Featureless classes or classes with small particle number were discarded. In total we were able to separate six principal classes and 21 substates were refined to resolutions ranging from 10-12 Å for the principal states and 13-18 Å for the substates depending on the particle number.

**Measurement of central stalk rotation**
Two approaches were chosen to measure the angle by which the central stalk rotates during the transition from one principal state to the next. In both approaches, the central stalk was segmented from each map in the first step and the central stalk map of the assigned principal state I (see Fig. 4) was subsequently fitted into each of the other segmented central stalk using the “fit inMap” function in ChimeraX (*11*). In the second approach, the rotation of the central stalk between individual principal states was measured by fitting the segmented central stalk map from each individual states into the map of the following state using the same command. Both strategies resulted in similar values with ± 1°. The angles in Fig.S5 are the mean values of both measurements.

**Nearest-neighbour analysis**
Nearest-neighbour analysis was performed using our own python scripts executed in a Jupyter notebook. The data was displayed using napari (*12*).

**Membrane segmentation and 3D representation**
Membranes were segmented using MemBrain v2 (*13*). 3D visualization of the ATP synthase dimer along the inner mitochondrial membrane was performed in ChimeraX (*11*) using the ArtiaX toolbox (*14*). All figures were prepared with ChimeraX.

**Model building**
To build the model into the consensus map, a single particle cryoEM structure of ATP synthase dimer from the same organism was used as a template (PDB:6rd4). Initially, the structure was docked in to the map using ChimeraX ‘fit in’ function. The model was inspected in *Coot* (*15*) and adjusted as necessary. A major large unassigned density, not present in the single-particle structure, originated from the N-

terminus of ASA3. The N-terminal ASA3 amphipathic and map helices (residue 25-83) were modelled using the homology model structure from *Chlamydomonas reinhardtii* (Uniprot code: A8JCE9). The sequences were then mutated using Coot's 'mutate residue range' function. Additionally, the loop between transmembrane helix 2 and 3 of subunit *a* (residues 205-218) was built starting with the Alphafold (*16*) model as a reference and refined it to the density (*16*). To globally refine the structure, ISOLDE was implemented with the use of strict distance and torsion restraints, which were released only for the newly modelled segments (*17*). Finally, the 'phenixRefineInput' command in ISOLDE was used to generate a file for *PHENIX* (*18*) with instructions to carry out only global minimization and B-factor (ADP) refinement.

A

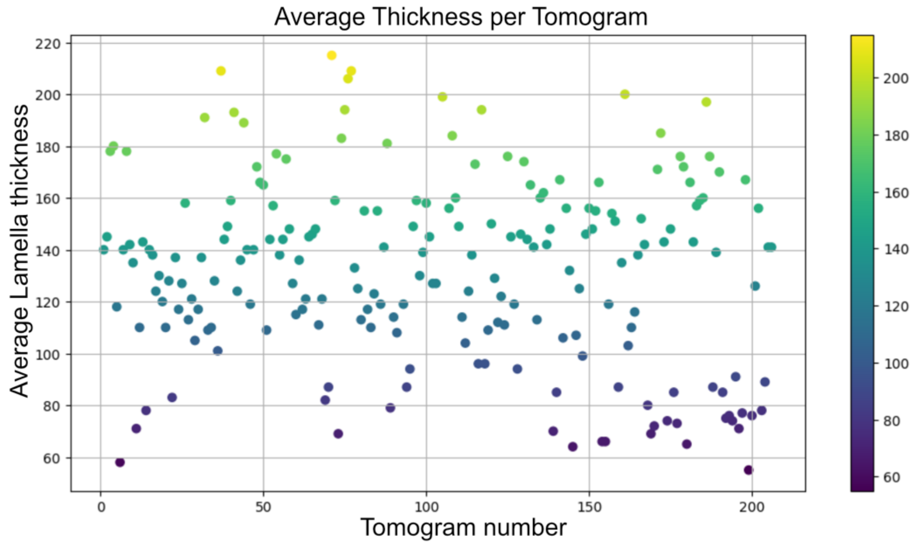
Average Thickness per Tomogram
Average Lamella thickness
Tomogram number

B

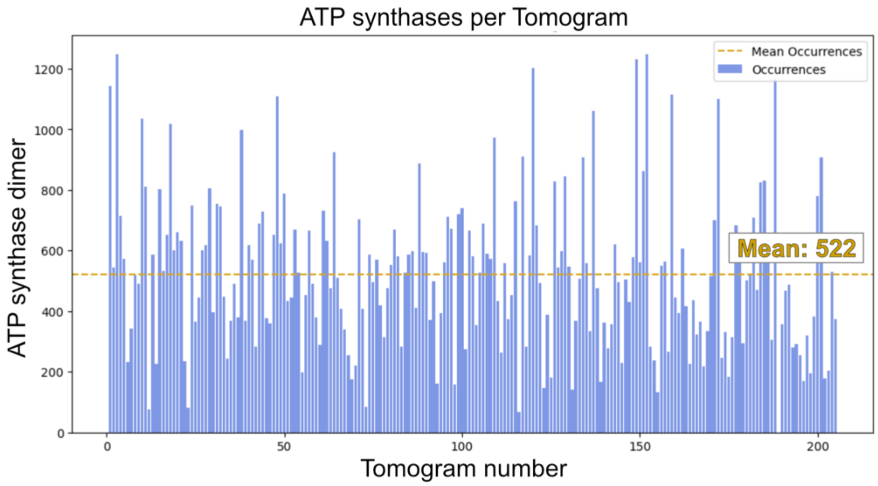
ATP synthases per Tomogram
Mean Occurrences
Occurrences
Mean: 522
ATP synthase dimer
Tomogram number

**Fig. S1**. **Statistics of lamella thickness and ATP synthase dimer per tomogram. (A)** Average thickness of lamellae cut by FIB milling used for tomographic data collection. The thickness of each tomogram was measured manually at three points. (**B)** Number of ATP synthase dimer volumes extracted from each tomogram, as indicated by the consensus dimer map. The figures reflect the quality of individual tomograms rather than the actual number of ATP synthases present in each tomographic volume.

#### Processing workflow I

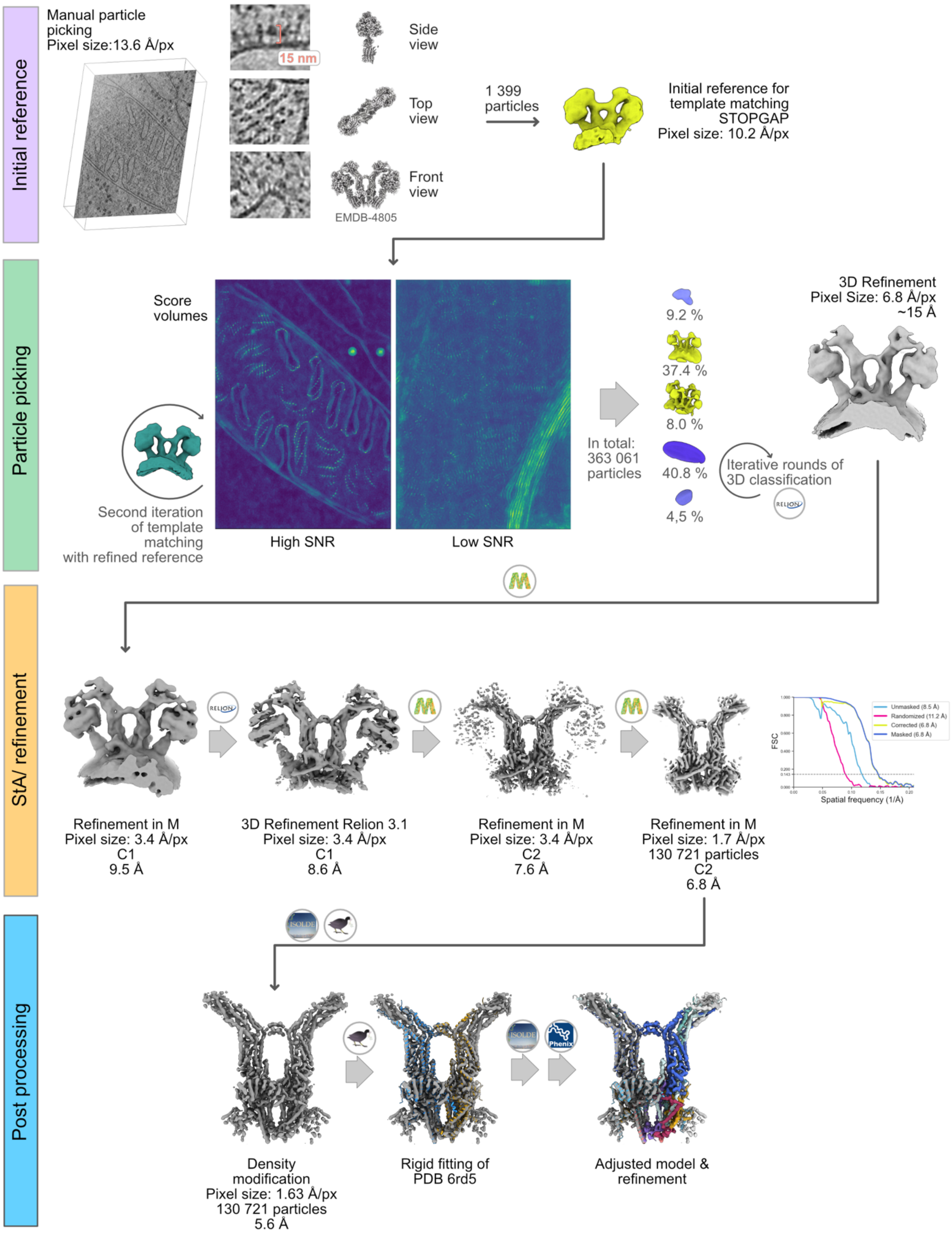

**Fig. S2: Workflow for subtomogram averaging (part 1).** Starting with the reconstructed tomograms, individual ATP synthase particles were visually identified by their characteristic shape using the published single-particle map (*19*) as a reference. An initial reference was generated from 1,399 particles. Template matching was performed with the program STOPGAP (*8*). The resulting score volumes differed widely with respect to signal-to-noise ratio (SNR). True positives were identified from the score volumes through their formation of long rows. Extensive false positive picks were observed along all kinds of membranes. False positive picks were sorted out by iterative rounds of 3D classification in Relion3.1 (*5*), yielding an initial consensus map of 15 Å resolution. The consensus map was further refined with the programs M(*20*) and Relion3.1(*5*). Focused refinement and masking of the peripheral stalk with imposed C2 symmetry produced a final map at a global resolution of 6.8 Å, with a local resolution up to 4.2 Å. Density modification with the *PHENIX* toolbox (*18*) improved the global resolution to 5.6 Å. The published 2.8 Å single-particle structure of the ATP synthase dimer (PDB: 6rd4) was fitted into the 5.6 Å density modified map and refined in Coot (*15*) and ISOLDE (*17*). Major adjustments to subunits ASA3 and subunit a were made in Coot. Final small-scale adjustments were made in ISOLDE and finally globally refined in *PHENIX* using the 6.8 Å consensus map using supplied distance-distance and torsion restraints from ISOLDE.

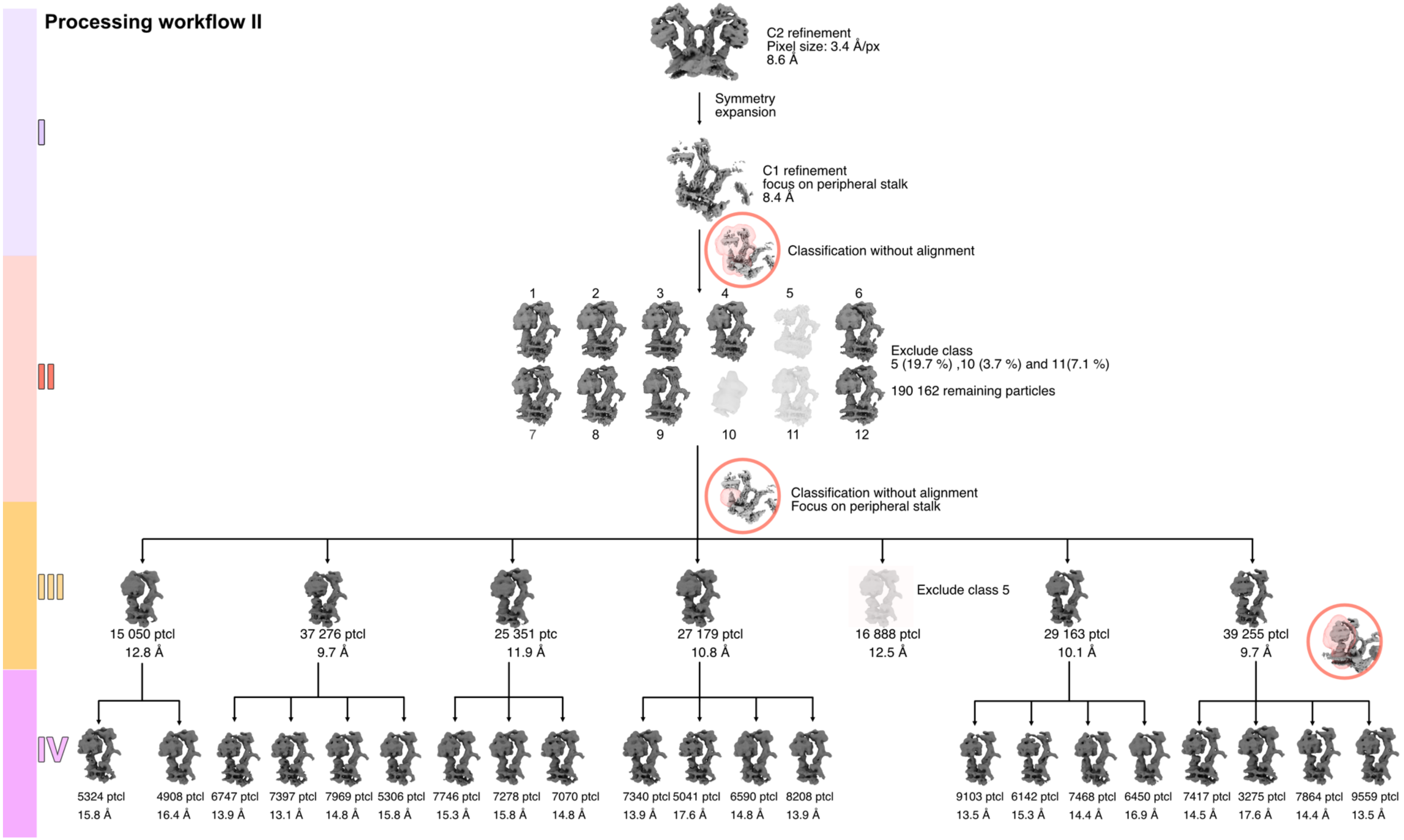

Processing workflow II
I
II
III
IV
C2 refinement
Pixel size: 3.4 Å/px
8.6 Å
Symmetry expansion
C1 refinement
focus on peripheral stalk
8.4 Å
Classification without alignment
1
2
3
4
5
6
7
8
9
10
11
12
Exclude class
5 (19.7 %) ,10 (3.7 %) and 11(7.1 %)
190 162 remaining particles
Classification without alignment
Focus on peripheral stalk
15 050 ptcl 12.8 Å
37 276 ptcl 9.7 Å
25 351 ptc 11.9 Å
27 179 ptcl 10.8 Å
16 888 ptcl 12.5 Å
Exclude class 5
29 163 ptcl 10.1 Å
39 255 ptcl 9.7 Å
5324 ptcl 15.8 Å
4908 ptcl 16.4 Å
6747 ptcl 13.9 Å
7397 ptcl 13.1 Å
7969 ptcl 14.8 Å
5306 ptcl 15.8 Å
7746 ptcl 15.3 Å
7278 ptcl 15.8 Å
7070 ptcl 14.8 Å
7340 ptcl 13.9 Å
5041 ptcl 17.6 Å
6590 ptcl 14.8 Å
8208 ptcl 13.9 Å
9103 ptcl 13.5 Å
6142 ptcl 15.3 Å
7468 ptcl 14.4 Å
6450 ptcl 16.9 Å
7417 ptcl 14.5 Å
3275 ptcl 17.6 Å
7864 ptcl 14.4 Å
9559 ptcl 13.5 Å

**Fig.S3. Workflow for subtomogram averaging (part 2).** Level I (lilac): Starting with the consensus map with imposed C2 symmetry, the symmetry was expanded, followed by focused refinement of the peripheral stalk in C1. Classification masking of the monomer including the peripheral stalk sorted out particles with insufficient signal in the $F_1F_o$ part of the protein, resulting in a total of 190,162 particles which were subjected to focused classification to dissect principal rotary states (Level I, red). The principal states were classified without alignment into seven classes using a focused mask around the lower portion of the asymmetric central stalk sitting above the c-ring (Level III, orange). A second round of focused classification masking the $F_1F_o$ part (excluding the peripheral stalk) was performed for each principal rotary state. This approach resulted in 2 to 4 subclasses per principal state (Level IV, pink), indicating differences in the position of the $F_1$ head around the central stalk axis. Close inspection suggested that there were more substates present in the data, however limited particle numbers prohibited a finer sampling of the rotational landscape. Featureless classes were discarded (not shown).

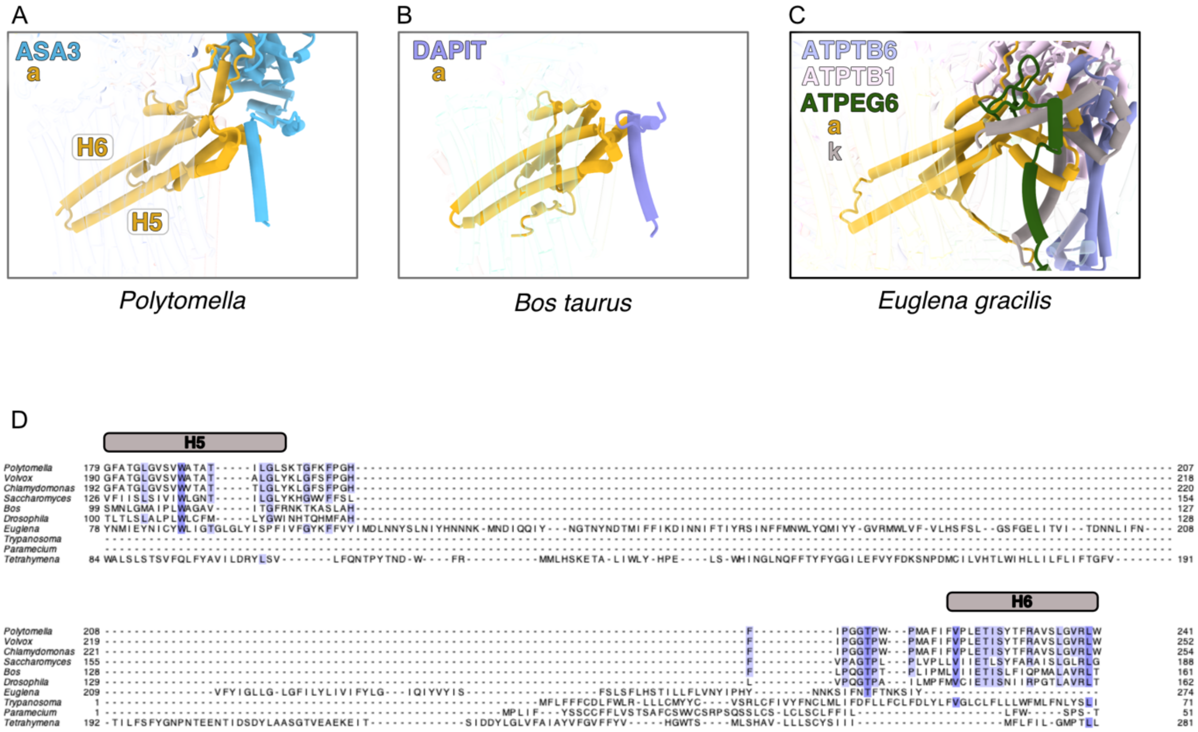
A
ASA3
a
H6
H5
Polytomella
B
DAPIT
a
Bos taurus
C
ATPTB6
ATPTB1
ATPEG6
a
k
Euglena gracilis
D
H5
H6
Polytomella
Volvox
Chlamydomonas
Saccharomyces
Bos
Drosophila
Euglena
Trypanosoma
Paramecium
Tetrahymena

**Fig.S4. The H5-H6 loop stabilizes subunit *a* in mitochondrial ATP synthases of different organisms**. The ASA3 loop and its interacting subunits are coloured and labelled for **(A)** *Polytomella* (*P. pringsheim*, type II ATP synthase), **(B)** *Bos taurus* (type I ATP synthase, 6ZQM) and **(C)**, *Euglena gracilis* (type IV ATP synthase, 6TDU). **(D)** Sequence alignment of ATP synthase from all four types highlight the sequence conservation of the H5-H6 loop in types I and II. ATP synthase type I: *Bos taurus, Drosophila melanogaster, Saccharomyces cerevisiae*, type II: *Polytomella pringsheim*, *Volvox carteri*, *Chlamydomonas reinhardtii*, type III: *Trypanosoma brucei*, *Tetrahymena thermophila*, type IV: *Euglena gracilis*, *Paramecium tetraurelia*.

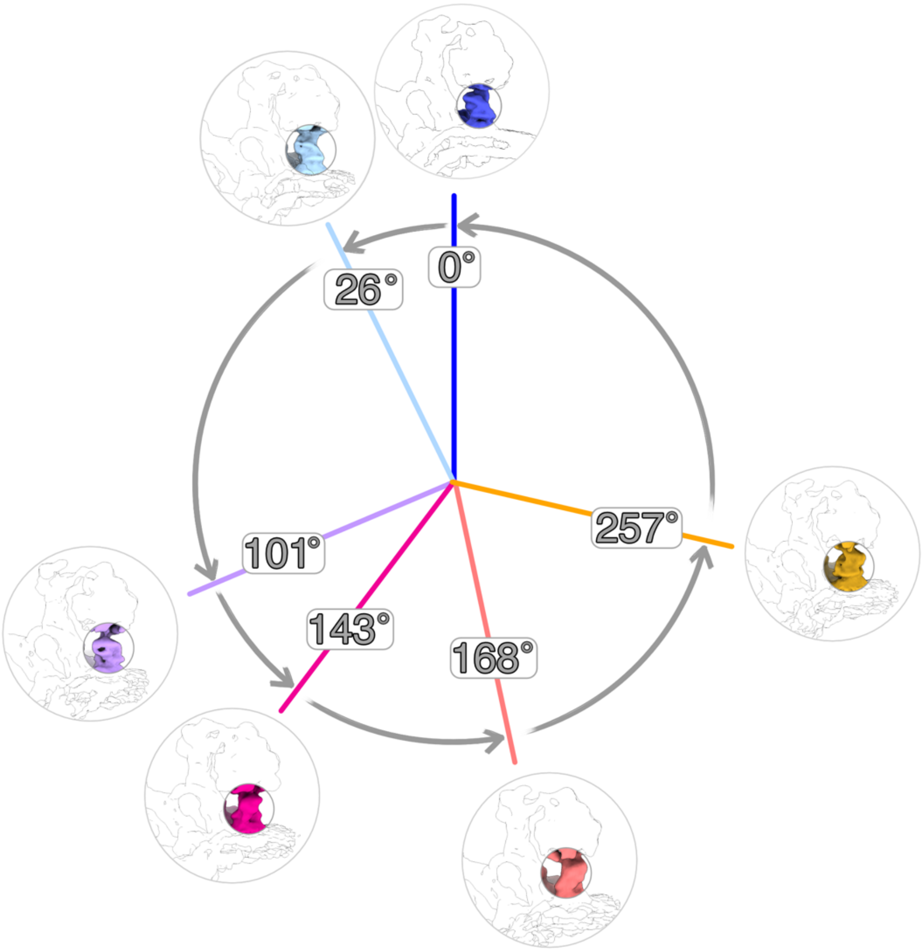
0°
26°
101°
143°
168°
257°

**Fig.S5. Rotation angles of the principal rotary states of the ATP synthase central stalk as observed in our *in situ* dataset.** 0° is assigned to the state that most closely corresponds to the primary rotation state 1 in the single-particle map (PDB:6rd9)(*19*). The anti-clockwise rotation of the central stalk was measured using the central stalk of state 1 as a reference. The rotation of the central stalk was measured by using the central stalk of the first rotation state as a reference and measuring the rotation required to fit it into each of the subsequent rotation states using the "fit inMap" command in ChimeraX (*11*). The angles were confirmed by measuring the rotation required to superimpose the central stalk for each state with the subsequent state. The angles differ from the 120-degree steps described in vitro, but represent potential further energy minima of the ATP synthase in the presence of a membrane potential *in situ*.

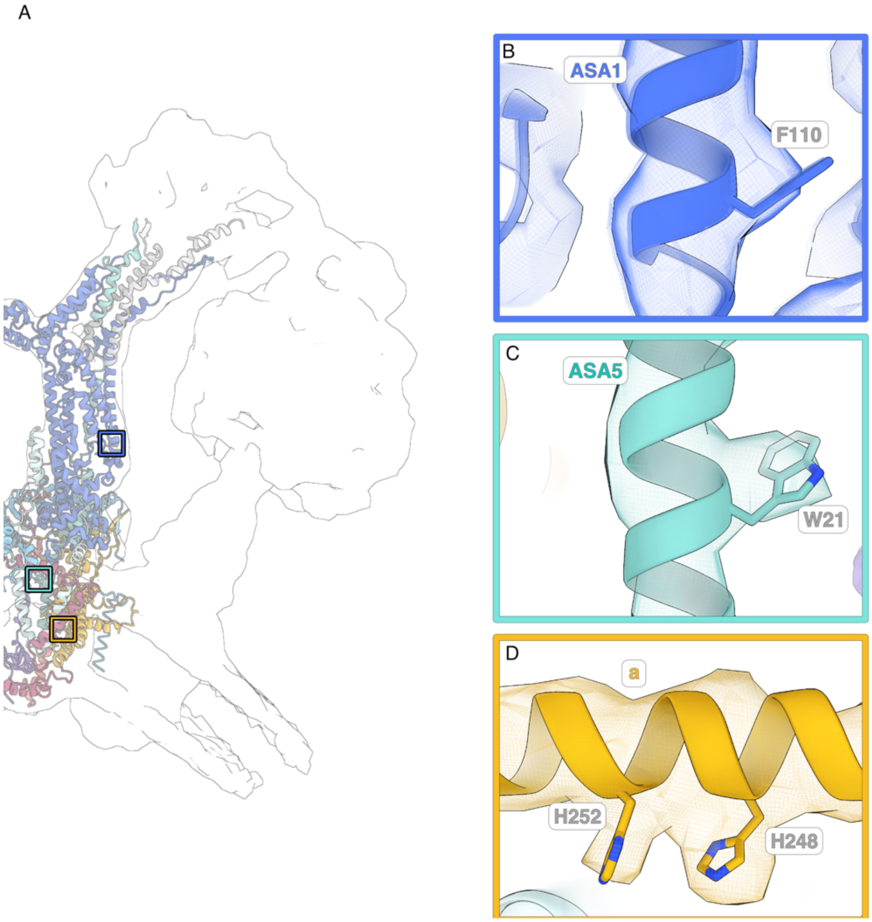
A
B
ASA1
F110
C
ASA5
W21
D
a
H252
H248

**Fig.S6. Docking the model (PDB:6rd4) to the *in situ* subtomogram average map of *Polytomella* ATP synthase. (A)** The resolution of the peripheral stalk was sufficient to see bulky aromatic side chains which were used as guide points for fitting the model into the map and ensuring the correct sequence register. (**B-D)** Closeups of representative areas (boxes in A) showing clear density of bulky aromatic side chains. B: Phenylalanine from ASA1 located at the matrix-facing side of the complex. **(C)** Tryptophan in ASA5 located at the core of the protein inside the membrane. **(D)** Two histidines coordinating a metal ion in the horizontal helix hairpin of the subunit *a* (*10*). The metal ion was not resolved in the subtomogram average map. Isosurface threshold: 0.1 for ASA1:F110 and a:H248,252 and 0.2 for ASA5:W21.

**Table S1. Summary of the peripheral stalk model of Polytomella ATP synthase**

| Subunit | Chain ID | Total residues | Modelled (%) |
|---|---|---|---|
| ASA10 | 0 | 82 | 2-82 (98.8) |
| ASA1 | 1 | 618 | 23-617 (96.3) |
| ASA3 | 3 | 325 | 25-321 (92.6) |
| ASA5 | 5 | 123 | 1-123 (100) |
| ASA6 | 6 | 151 | 18-151(88.7) |
| ASA7 | 7 | 190 | 79-190 (58.9) |
| ASA8 | 8 | 89 | 2-89 (98.9) |
| ASA9 | 9 | 97 | 1-97 (100%) |
| a | M | 327 | 95-324 (70.3) |

**Movie S1**

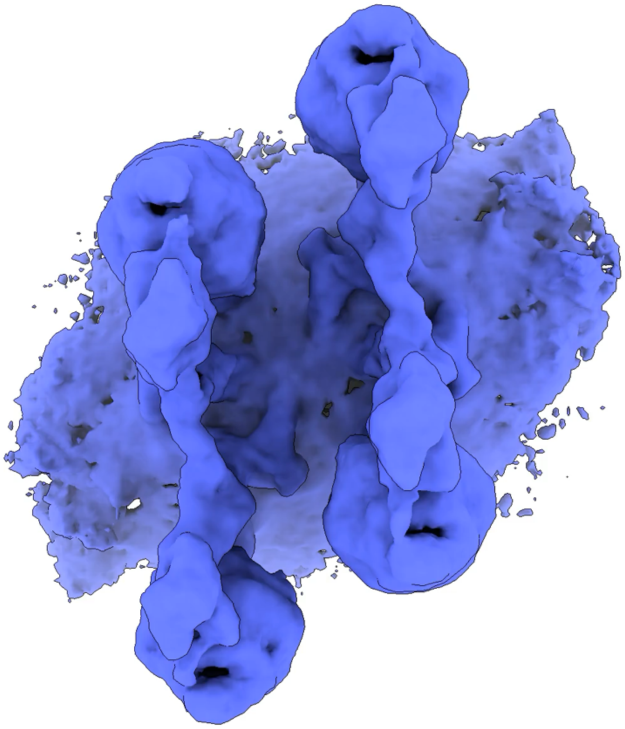

Morph between different classes resulting from the 3D classification of ATP synthase tetramers. Although the distance between the tetramers is precisely defined, individual dimers remain flexible in relation to each other and can thus adapt to the local crista morphology.
